# Transcriptomic analysis and high throughput functional characterization of human induced pluripotent stem cell derived sensory neurons

**DOI:** 10.1101/2024.08.23.607310

**Authors:** Vincent Truong, Jackson Brougher, Tim Strassmaier, Irene Lu, Dale George, Theodore J. Price, Alison Obergrussberger, Aaron Randolph, Rodolfo J. Haedo, Niels Fertig, Patrick Walsh

## Abstract

Peripheral sensory neurons are a primary effector in pain neurotransmission, and have become a useful cellular model for the study of pain. While rodent tissue has historically served as a source of these neurons, it has become increasingly clear that pain mechanisms in rodents and humans are substantially divergent. Sensory neurons harvested from cadaveric human tissue serve as a superior translational model for studying pain mechanisms, however their relative paucity limits their widespread utility. Theoretically, sensory neurons manufactured from human pluripotent stem cells (hPSCs) could help bridge this translational gap given their relative abundance and potential similarity to primary human tissue. However, hPSC-derived sensory neurons manufactured with the most common methodologies correlate poorly to human tissue both transcriptionally and functionally. In the present work, we compare a novel population of hPSC-derived sensory neurons to previously published datasets and find this novel population to more closely resemble human primary dorsal root ganglia transcriptionally. Furthermore, we evaluate the heterogeneity of this novel population via single nucleus RNA sequencing and find it resembles specific nociceptor and mechanoreceptor subsets found in vivo. Finally, we assay the functionality of this population with high throughput automated patch clamp electrophysiology with respect to voltage-gated sodium (Na_v_) and potassium channels (K_v_), and ligand-gated ionotropic GABA and P2X receptors. Overall, we find this population of hPSC-derived sensory neurons to be of relatively high fidelity, and suitable for interrogating numerous potential pain targets on a fully humanized platform.

## Introduction

Pain is a complex and debilitating condition that will affect nearly every living human throughout their lifetime. Despite significant advances in our understanding of the molecular and cellular mechanisms underlying pain, the development of effective pain therapeutics that are devoid of major side-effects or toxicities has been challenging. Part of this difficulty stems from the limited availability of human models for both pain drug discovery and fundamental, mechanism-based research [1]. Sensory neurons, a key component of the peripheral nervous system, play a crucial role in conveying nociceptive signals from the body to the spinal cord and brain where pain is ultimately perceived [2]. Traditional preclinical models predominantly utilize primary rodent dorsal root ganglion (DRG) cells, the overexpression of ion channel targets in human embryonic kidney (HEK) or Chinese hamster ovary (CHO) cell lines, or ideally human DRG – which is a scarce resource for most laboratories [1]. Human induced pluripotent stem cell (hiPSC)-derived sensory neurons are a scalable and pertinent alternative, offering a viable and efficient approach for modeling pain in high-throughput drug discovery efforts [3].

Multiple human iPSC sensory neuron protocols exist in the literature based on two traditional methods. The first are directed differentiation methods replicating nervous system development using the classical dual SMAD inhibition based neural induction [4] or nocispheres [5], patterning to a neural crest intermediate, and then terminal differentiation to sensory neurons with varying levels of efficiencies [6–9]. Alternatively, there are protocols for the overexpression of cell-fate determining transcription factor programming to generate mechanoreceptors [10] or nociceptors [11]. A recent, novel method of directed differentiation through an intermediate primal ectoderm population [12] has been shown to reproducibly generate sensory neurons (Anatomic RealDRG) [13, 14] where voltage gated sodium [15, 16] and Piezo2 [17, 18] channels have been functionally validated – along with their utility for optimization of cryopreservation [13] and numerous other applications [14, 19–23].

In the studies described here, we have thoroughly characterized RealDRG neurons throughout their maturation using RNA sequencing, as well as automated patch clamp electrophysiology (APC). This was done with the goal of understanding the utility of these neurons for drug screens and mechanistic studies, and to create a thorough resource for researchers who would use these neurons for these or other purposes. We have bulk RNA sequenced RealDRG through four weeks of maturation and compared the expression to both primary hDRG [24, 25] and other hiPSC-derived sensory neurons. Using single nuclei sequencing, we also look at how maturation develops through the first two weeks to determine what sensory neuron sub-type RealDRG are most similar to in hDRG. Finally, we sought to functionally validate a panel of expressed targets utilizing high-throughput APC electrophysiology.

## Results

### 1. hiPSC-derived sensory neuron principal component analysis and comparison to hDRG

To determine how the bulk transcriptome of hDRG and hiPSC-derived sensory neuron populations compare, a number of published datasets [6, 8, 10, 11] were compiled from GEO and principal component analysis (PCA) was performed to explore the time course differentiation and maturation of hiPSC sensory neurons generated using various protocols. These datasets were compared to hiPSC lines [6, 8, 10], bulk sequenced hDRG, which contains a mix of neuronal subtypes, glia and other cells [10, 24], and the novel differentiation method currently disclosed [26] **(Figure 1)**. Initially, unbiased analysis using the top 2000 variable genes failed to display any distinct trends, likely because of major differences in the total number of cell types found in each of the bulk RNA sequencing datasets. Therefore, we focused the analysis on a previously defined panel of 173 hDRG-specific genes conserved between mice and human [24]. This more focused analysis showed different molecular signatures between the cell populations analyzed and that no hiPSC derived sensory neuron population overlapped exactly with primary hDRG, though some datasets were more similar than others.

**Figure 1.**
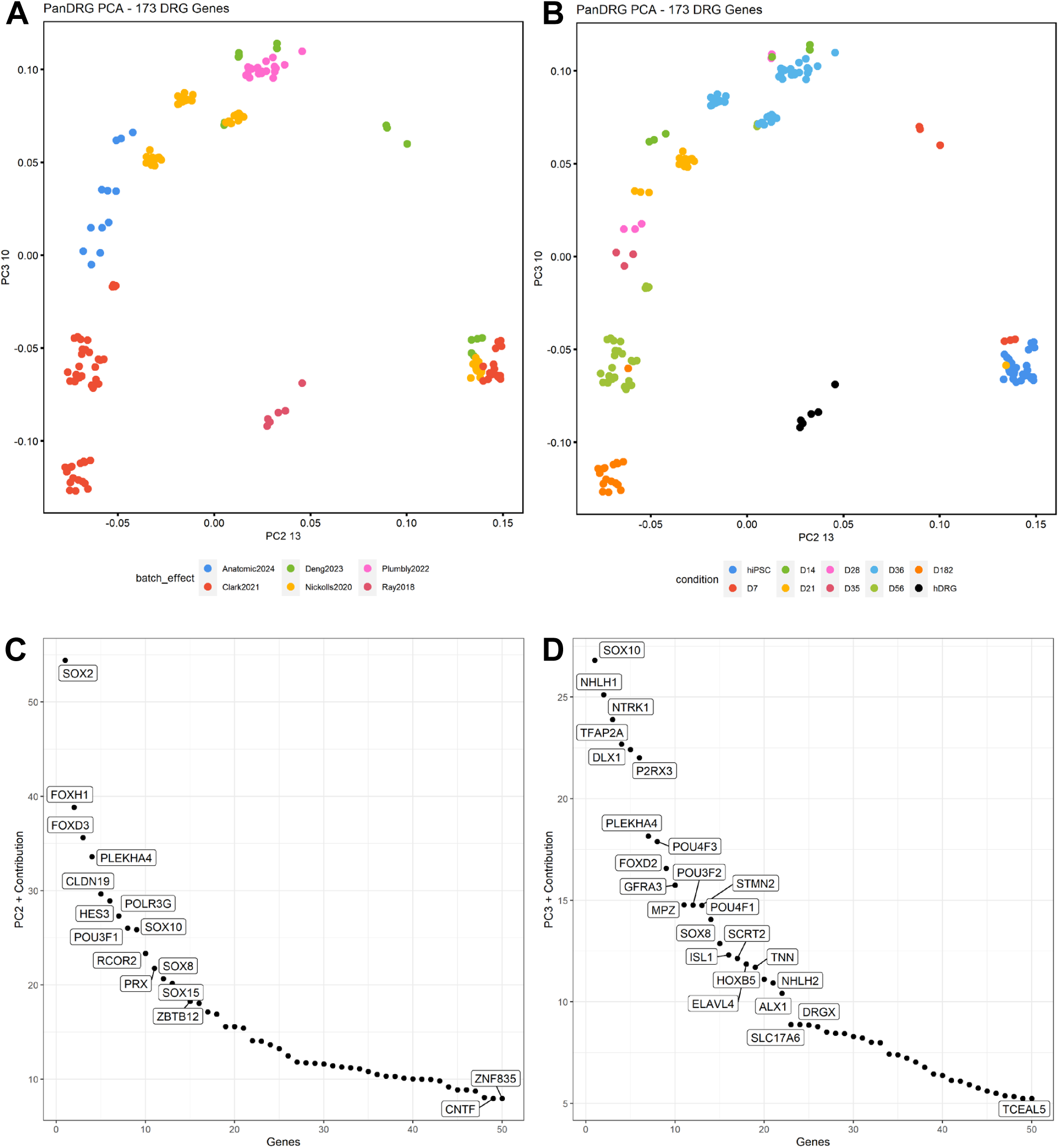
Transcriptomic analysis using a panel of 173 sensory neuron specific genes with data sets including primary human DRG, hiPSC-derived sensory neurons, and human induced pluripotent stem cells show differentiation and maturation trajectories. **A)** PCA plot of multiple RNA-seq data sets show distinct molecular signatures by protocol. **B)** PCA plot of multiple RNA-seq data sets show distinct molecular signatures at various timepoints. **C)** A list of genes that contribute to the variance of principal component 2 (PC2). **D)** A list of genes that contribute to the variance of principal component 3 (PC3).

Since the datasets included contain multiple cell types upon a differentiation continuum, including hiPSCs, neural crest, sensory neurons over differing maturation timeframes, as well as adult human primary DRG; we first explored principal component space to determine which principal components reflected a differentiation trajectory to be able to describe the relative maturation and identity of these populations. We found that while principal component 1 was primarily populated by non-specific housekeeping genes, principal components 2 and 3 provided a rational developmental progression from hPSC to ahDRG (**Figure 1**). The lower right corners of each plot in **Figure 1** are occupied by hPSCs, indicating the embryologically least mature cell state. Deng et al. provide valuable embryological context by annotating the differentiation stages in their chronological time course. They describe their ’day 7 population’ as ’neural ectoderm,’ which are a direct embryological descendant from hPSCs. By ’day 14,’ this population differentiates into ’neural crest,’ which further matures into the ’nociceptor’ population by ’day 56.’

To validate the hypothesis that PC2/PC3 describes an embryological trajectory, we further decomposed each principal component PC2 and PC3 into the genes from which they’re comprised (**Figure 1C, 1D**). Contributors to PC2 and PC3 include known transcription factors corresponding to neurodevelopment, such as SOX2, POU3F1, and SOX10 in PC2; and DLX2, POU4F3, and SOX8 in PC3. Additional genes comprising PC2 and PC3 are related to sensory function, such as nociceptive channels NTRK1 and P2RX3; indicating the analysis to be valid.

Given these landmarks, we can observe in PC2/PC3 that embryological time rotates counterclockwise from the lower right hand quadrant of the graphs in **Figure 1A and 1B**, with the embryologically most mature populations eventually occupying the bottom-left quadrant of the PC2/PC3 plane. Using this map, we can evaluate which populations included are embryologically most mature. The embryological maturation order, therefore, becomes hPSC, Deng days in vitro (DIV) 7, Plumbly DIV 36, Deng DIV 14, Deng DIV 21, Deng DIV 56, Nickolls DIV 36, Anatomic DIV 14, Nickolls DIV 21, Anatomic DIV 21, Anatomic DIV 28, Anatomic DIV 35, Clark DIV 56, Clark DIV 182. As should be obvious from this analysis, the chronology – or DIV— doesn’t necessarily predict embryological maturation. Differentiation protocols can be of more advanced embryological age despite a younger chronological age—and vice-versa—reflecting differences between differentiation methods.

To expand upon this point, it appears that differentiation methods reliant upon transcription factor over-expression, the Nickolls and Plumbly datasets, remain embryologically immature and reminiscent of neural crest cells despite an advanced chronological age, which is surprising due to these cells displaying an obvious neuronal phenotype in culture. On the other hand, directed differentiation methods reliant upon embryonic morphogens (Anatomic, Clark, Deng) display the potential to embryologically mature with greater time in vitro. This embryological maturation timescale differs chronologically depending on the protocol, with each protocol having its own maturation timescale. Clark et al appears to embryologically mature chronologically faster than Deng, and Anatomic faster than Clark and Deng.

Notably, no hiPSC-derived population fully overlaps with hDRG in PC space. This is likely due to the relative heterogeneity of the hDRG, as this dataset was from bulk-sequenced whole DRG, which includes numerous non-sensory neuron cell types such as glia, fibroblasts, and immune cells.

### 2. Ion channel-, neuropeptide-, and receptor-based comparison of iPSC sensory neurons datasets to native hDRG expression

While the PCA analysis described above gives insight into similarities in transcriptome based on DRG-specific gene expression, pharmacological targets expressed in hDRG including ion channels, neuropeptides, GPCRs, and cell adhesion molecules (CAMs) [24] provided further insight on what differentiates the hiPSC-derived sensory neuron populations (**Figure 2**). Ion channels play a critical role in DRG neuron physiology. Among these channels are voltage gated sodium and calcium channels, transient receptor potential (TRP) channels, PIEZO channels, and purinergic channels that are all involved in nociceptive signal transduction in DRG neurons. In general, ectoderm-derived hiPSC sensory neurons expressed the most ion channels out of all the hiPSC sensory neuron populations (**Figure 2A**). Clark et al. also had expression of a number of ion channels – perhaps due to the longer time of maturation in comparison to all protocols or the differentiation method. The voltage gated sodium channels SCN9A (Na_V_1.7) and SCN10A (Na_V_1.8) are common pain targets due to known genetic mutations which either cause gain of function (erythromyalgia) or loss of function (pain agnosia). RealDRG SCN10A expression was highest compared to all protocols and SCN9A expression was similar to hDRG samples. The Clark et al. protocol had highest expression of SCN9A and SCN11A, which is integral to the fine-tuning of neuronal excitability. The full list of voltage gated sodium channels can be seen in **Supplementary Figure 1A-B**. The TRPV1 receptor on C polymodal nociceptors and Aδ mechanoreceptors that are activated by noxious heat, capsaicin leading to burning pain or itch [27] is expressed at the highest level in the Clark et al. protocol and second highest in the RealDRG population. Notably, no hiPSC-derived sensory neuron expresses TRPV1 to the level of primary hDRG (**Supplementary Figure 1C-D)**, meaning hDRG far out-express their hiPSC-derived counterparts given sensory neuron transcripts account for only a fraction of the RNA sequenced in the hDRG samples given their heterogeneity. Interestingly of all the ion channels, Plumby et. al had a high level of expression of TRPA1, a common pain drug discovery target [28] involved with the detection of noxious stimuli including mustard oil, and wasabi. TRPA1 is also expressed in non-neuronal cell types involved with inflammation [29] including neural crest derived melanocytes which could be a contaminant in the culture [30]. A number of other pain targets including voltage gated potassium channels (KCNQ2/3, KCNQ4, and KCNMA1), the mechanosensitive Piezo2 channel, GABA receptors (GABRA2, GABRA3, GABRB3), the acid sensing channel ASIC1, and purinergic channels (P2RX3, P2RX5) were all highly expressed in RealDRG. In short, a large number of ion channels are expressed in RealDRG that could potentially be used as analgesic drug target screening applications [15, 17].

**Figure 2.**
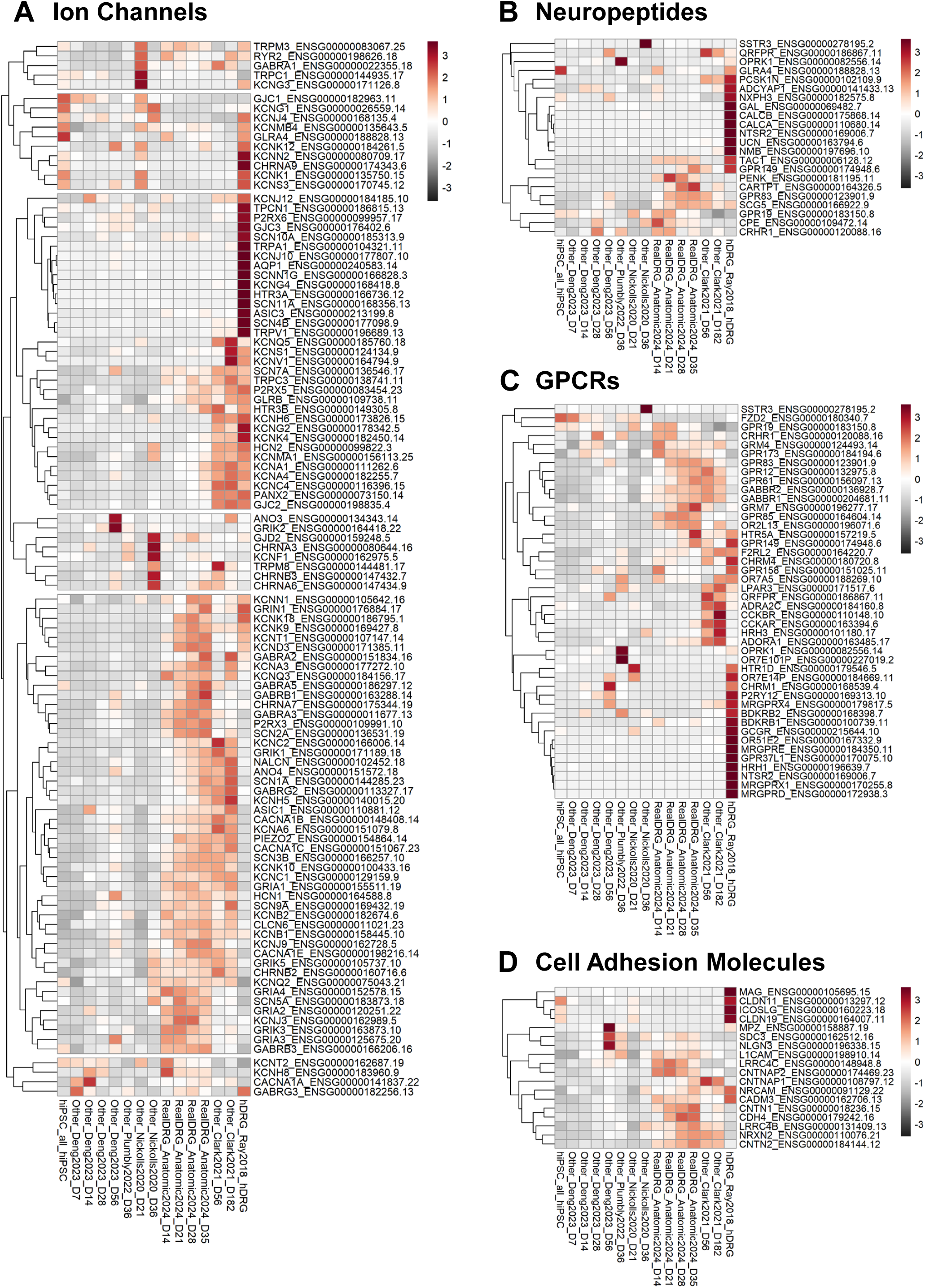
Heatmap analysis showing sensory neuron specific ion channel, neuropeptides, GPCRs, and cell adhesion molecules in primary human DRG, hiPSC-derived sensory neurons and hiPSCs. Expression of **A)** Ion Channels **B)** Neuropeptides **C)** GPCRs and **D)** Cell adhesion molecules.

Neuropeptides are often used as markers of subsets of sensory neurons and regulate signaling from DRG neurons to the vasculature and immune cells as well as to dorsal horn neurons in the spinal cord. RealDRG expressed more neuropeptides compared to the other hiPSC-derived sensory neurons, though there were a number of genes that were not expressed at the level seen in hDRG (**Figure 2B**). No hiPSC-derived sensory neuron population had expression levels of CALCA and CALCB similar to hDRG, which encode for the calcitonin gene-related peptide (CGRP) neuropeptide - a key migraine pain target using monoclonal antibodies and small molecule receptor inhibitors [31–36]. RealDRG expressed the highest level of TAC1, (tachykinin 1) a gene that encodes for the precursor protein of substance P. They also expressed PENK (Proenkephalin) a key precursor to enkephalins, which are endogenous opioid peptides that bind to mu-opioid receptors (MORs) and produce analgesia [37, 38].

At least 40 members of the approximately 400 non-olfactory G protein coupled receptors (GPCRs) in the human genome are targets for the regulation of pain [39, 40]. RealDRG as well as the Clark et al. sensory neurons expressed the most GPCRs that were also found in hDRG (**Figure 2C**). RealDRG expressed GRM7 (Glutamate Receptor, Metabotropic 7) and HTR4A (5-Hydroxytryptamine Receptor 4A) which both play a pivotal role in modulating neurotransmitter signaling and sensory processing. OPRK1, or kappa-opioid receptor 1, is a gene that encodes for a protein that is involved in the regulation of pain sensation was also expressed within this population. Clark et al. highly expressed MRGPRX4 and MRGPRE, genes that encode Mas-related G protein-coupled receptors (Mrgprs) that are involved in the regulation of pain and itch sensations.

CAMs are involved with inflammatory and cell signaling processes and are increasingly viewed as pharmacological targets [41]. RealDRG express the most CAM pharmacological targets like CNTN1, NLGN3, NRCAM, and LRRC4C (**Figure 2D**). Deng et. al and Plumby et. al have expression of myelin protein zero (MPZ) and myelin-associated glycoprotein (MAG), which are related to myelin formation and maintenance in the nervous system indicating potential Schwann cell contaminants that could originate from the common neural crest progenitor. Together, RealDRG and Clark et al. sensory neurons expressed the most ion channels, neuropeptides, GPCRs, and cell adhesion molecules.

### 3. Single nucleus data with differentiation path framework from Day 0-14

To assess homogeneity of maturing sensory neurons and track expression of key nociceptor markers such as NTRK2, SCN9A, and SCN10A over time we performed single nuclei RNA sequencing on RealDRG cells cultured to DIV0, DIV7, DIV10, and DIV14 post-differentiation (**Figure 3A**). 26,733 nuclei passed best practice quality control, with an average of 2,719 transcripts and 1,610 genes detected per nuclei. Leiden clustering detected 11 distinct clusters across the 4 timepoints. DIV0 and DIV14 cells predominantly exclusively populate clusters 5 and 2, respectively; whereas DIV7 and DIV10 are represented across clusters 1 and 3, and 0, 4, 6, and 7, respectively. This seems to suggest an initial divergence from a homogeneous population into heterogeneity, followed by a convergence back into a homogenous population; possibly suggestive of selective events or resolution of transient intermediate fates. This is supported by genes represented in the terminal DIV0 and DIV14 timepoints, whose clusters 5 and 2 respectively are populated by neural crest genes (*PAX3, SOX5, TFAP2B*) followed by neuronal markers (*KCNAB1, KCNMB2, NAV3*) (**Figure 3B**). Notably, sample populations show relatively low levels of overlap across DIV, with the exception of Leiden clusters 7 and 8. The top five marker genes for cluster 7, *SNHG14, LRRC4C, NRG3, LINGO2*, and *RYR3*, suggest this cluster may represent cells at an intermediate stage of differentiation towards a nociceptive phenotype. The presence of *RYR3*, a calcium channel involved in neuronal signaling, and *NRG3*, which is implicated in the development of peripheral nervous system, could indicate these cells are in the process of developing nociceptive functionalities, potentially of the C-fiber subtype. Conversely, the top five marker genes for cluster 8, *NEFL, MAP1B, NEFM, LINK-PINT*, and *7SK*, indicate a possible Aδ-fiber subtype.

**Figure 3.**
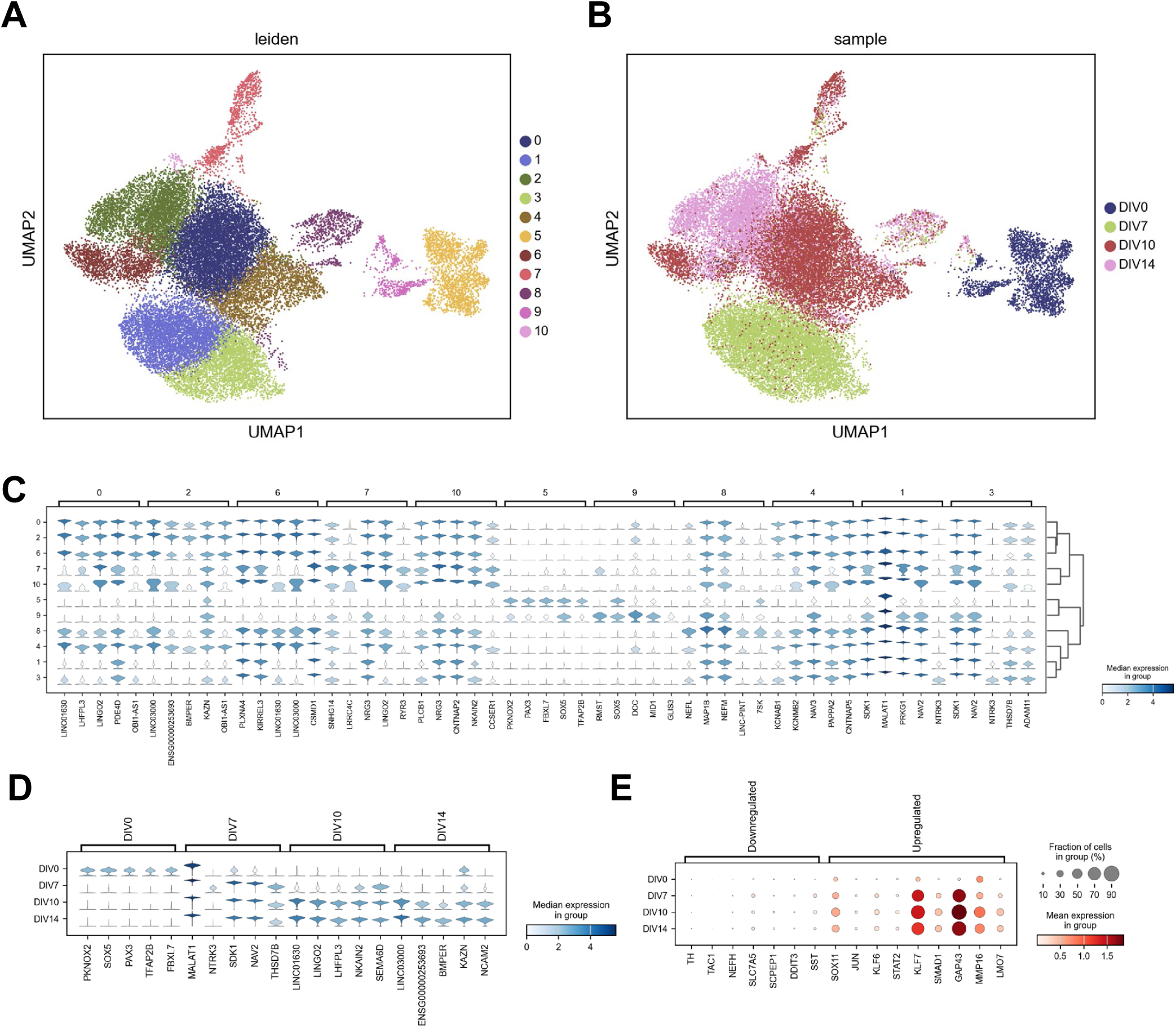
Single nucleus RNA sequencing of 26,733 sensory neurons at DIV0, DIV7, DIV10, and DIV14 post-differentiation reveals distinct transcriptomic signatures across time. **A)** UMAP plot showing 11 distinct clusters generated by Scanpy. **B)** UMAP plot showing distribution of individual sample timepoints. **C)** Stacked violin plot of top five marker genes for each individual Leiden cluster. **D)** Stacked violin plot of the top five marker genes for each individual timepoint. **E)** Dotplot showing mature sensory neurons resemble a neuropathic phenotype.

Next, we examined the top differentially expressed genes across samples (**Figure 3C**). DIV0 cells represented by cluster 5 seem to be in a neural crest or early sensory neuron state. *PKNOX2*, *SOX5*, and *PAX3* are transcription factors that play essential roles in early neural crest development and differentiation. *TFAP2B* is also involved in neural crest cell formation and its expression is a hallmark of early neural crest cells. *FBXL7* has a less well-known role in neural development, but its presence may suggest an active cellular state with ongoing protein turnover and differentiation. At DIV7, the cells appear to be transitioning towards a more mature sensory neuron phenotype. The expression of *NTRK3* (also known as TrkC), a receptor for neurotrophin-3, is indicative of differentiating sensory neurons [42] or indicative of low threshold mechanoreceptor neurons. Additionally, *NAV2* and *SDK1* are involved in axon guidance and synapse formation, suggesting ongoing maturation and connectivity of these cells. *MALAT1* is a long non-coding RNA known for its role in regulating gene expression, and its specific role here may be to facilitate this transition phase. By DIV10, the iPSC-derived cells seem to be further maturing into sensory neurons. *LINGO2* is involved in neuronal survival and regeneration, which may indicate ongoing maturation of these neurons. *SEMA6D* is involved in axon guidance, which further suggests maturation and connectivity. *LHFPL3* and *NKAIN2*, while less studied, have been associated with neuronal activity and might indicate functional maturation of these cells. By DIV14, the cells likely represent mature sensory neurons. *BMPER* is involved in the BMP signaling pathway, which is known to play a role in neuronal differentiation and maturation. *NCAM2* plays a role in neuron-neuron adhesion and in synapse formation, indicating that these neurons are likely forming synaptic connections. *KAZN’s* role in sensory neurons is less clear, but it is generally involved in cellular homeostasis.

Finally, we examined genes found in specific populations of nociceptors and also genes implicated in neuropathic pain or nerve injury (**Figure 3D**). The decreased expression of *TAC1*, *NEFH*, *SLC7A5*, *SCPEP1*, *DDIT3*, and *SST* suggests a potential lack of representation of neurons that express these markers in hDRG. On the other hand, the upregulation of genes like *SOX11*, *KLF7*, *GAP43*, *MMP16*, and *LMO7* implies an active state of neuronal regeneration or plasticity that is consistent with what is seen with nerve injury in animal models.

### 4. Single nucleus data with focus on nociceptor subtypes

We next chose to look at the previously examined expression of Ion Channels, Neuropeptides, GPCRs, and Cell Adhesion Molecules across identified Leiden clusters to determine the homogeneity of the RealDRG across maturation (**Figure 4**). Largely, differences in ion channel expression across Leiden clusters followed the separation of samples across clusters, such as the expression in cluster 5 which is predominantly made up of DIV0 nuclei (**Figure 4A**). The ubiquitous expression of *CPE* (Carboxypeptidase E) across all mature clusters from DIV7 onwards suggests a sustained requirement for neuropeptide processing in these differentiated neurons, as *CPE* is instrumental in the biosynthesis of peptide neurotransmitters and hormones (**Figure 4B**). Compared to total RNA sequencing, single nuclei sequencing detected abundant levels of relatively few GPCRs, likely due to the lower sequencing depth in the single nucleus experiments (**Figure 4C**). Notably, these do include *GABBR1*, *GABBR2*, *GPR173*, *GPR158*, and *GRM4*. The relatively high expression of these GABA receptors indicates expression of these targets for the use of positive controls for GPCR signaling in RealDRG. Further, deorphanizing of *GPR173* and *GRP158* may reveal unique targets for further manipulating cell excitability or explain previously unidentified drivers of signaling in hDRG. Lastly, single nuclei RNA sequencing detected relatively robust levels of cell adhesion molecules (**Figure 4D**). Together, these data indicate that across maturation, RealDRG represents a relatively homogenous group in their expression of ion channels and receptors.

**Figure 4.**
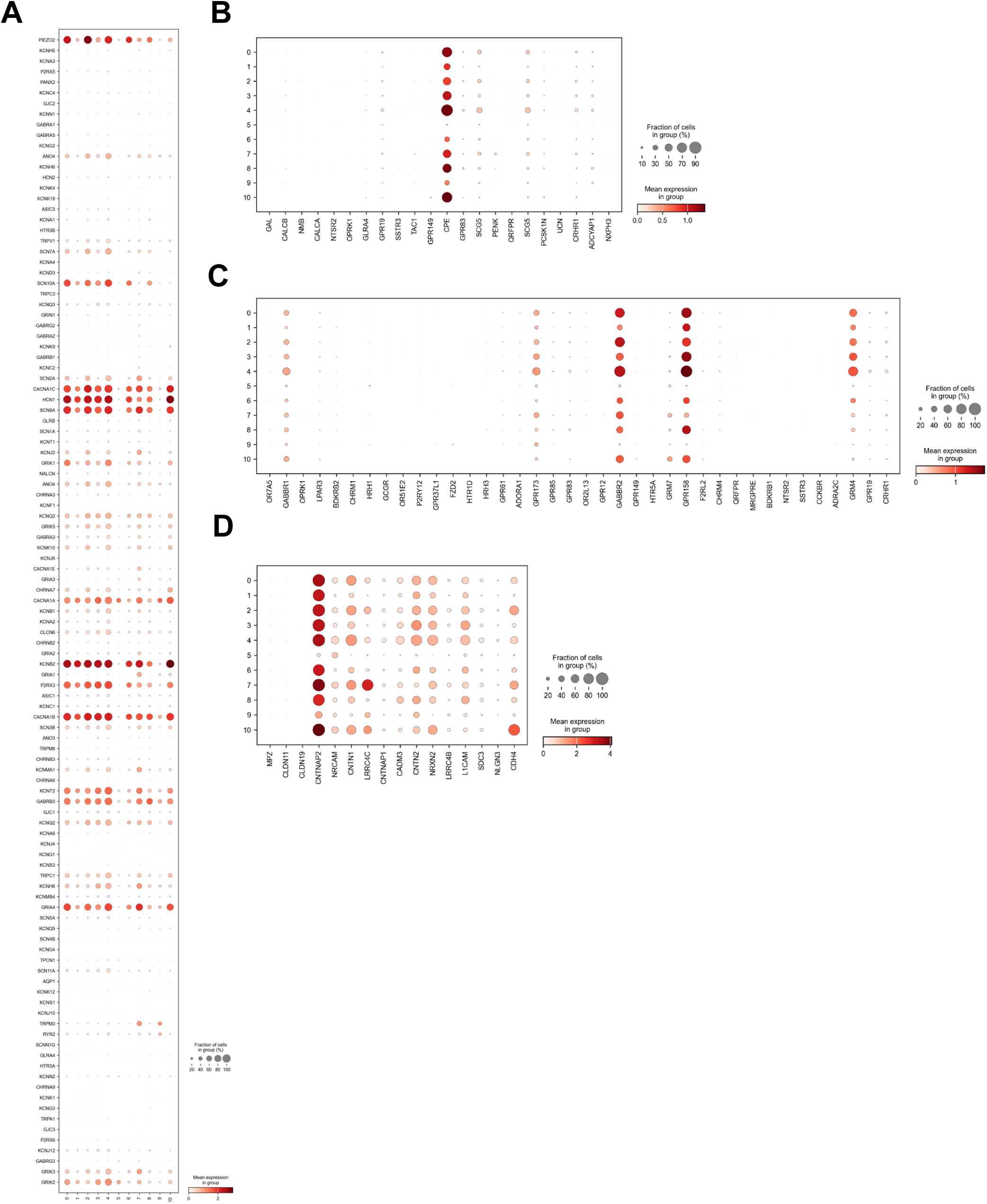
Dotplot analysis showing sensory neuron specific ion channel, neuropeptides, GPCRs, and cell adhesion molecules in single nucleus RNA sequencing Leiden clusters. **A)** Ion Channels **B)** Neuropeptides **C)** GPCRs **D)** Cell adhesion molecules

### 5. Comparison of single nuclei RNA sequencing cluster 2 to known nociceptive subtype marker genes

We next sought to understand if terminally differentiated RealDRG sensory neurons represented distinct nociceptor subtypes. First, we integrated a previously generated Visium dataset with proper batch correction using the *bbknn* function in Scanpy and performed a dendrogram clustering of nociceptor subtypes with the samples across time (**Figure 5A**). Doing so did not reveal any distinct integration of RealDRG samples with nociceptor subtypes. Next, we examined expression of key differentially expressed genes from nociceptor subtype clusters as previously performed with total RNA sequencing. Most notably, RealDRG express a range of Aδ**-**LTMR genes (KCNAB1, PIEZO2, ROBO2), Aδ**-**HTMR genes (SCN10A, MCTP1, NGFR), and select genes found in various nociceptor populations (PLXNA2, NTRK1, ATP2B4, KCNIP4) (**Figure 5B**). Given that the monolithic population found in cluster 2 – representing the most chronologically aged state in our dataset— appears to encompass multiple discrete populations found in the adult, we expect these neurons have yet to resolve their fate at this time point and possibly require additional maturation to determine their eventual subtypes. Regardless, it could be said that these immature neurons possess a dual-identity reminiscent of both mechanoreceptors and nociceptors. Sequencing of later timepoints would inform on whether a fate decision is eventually made to be either nociceptive, mechanoreceptive, or a mixed population following a divergence.

**Figure 5.**
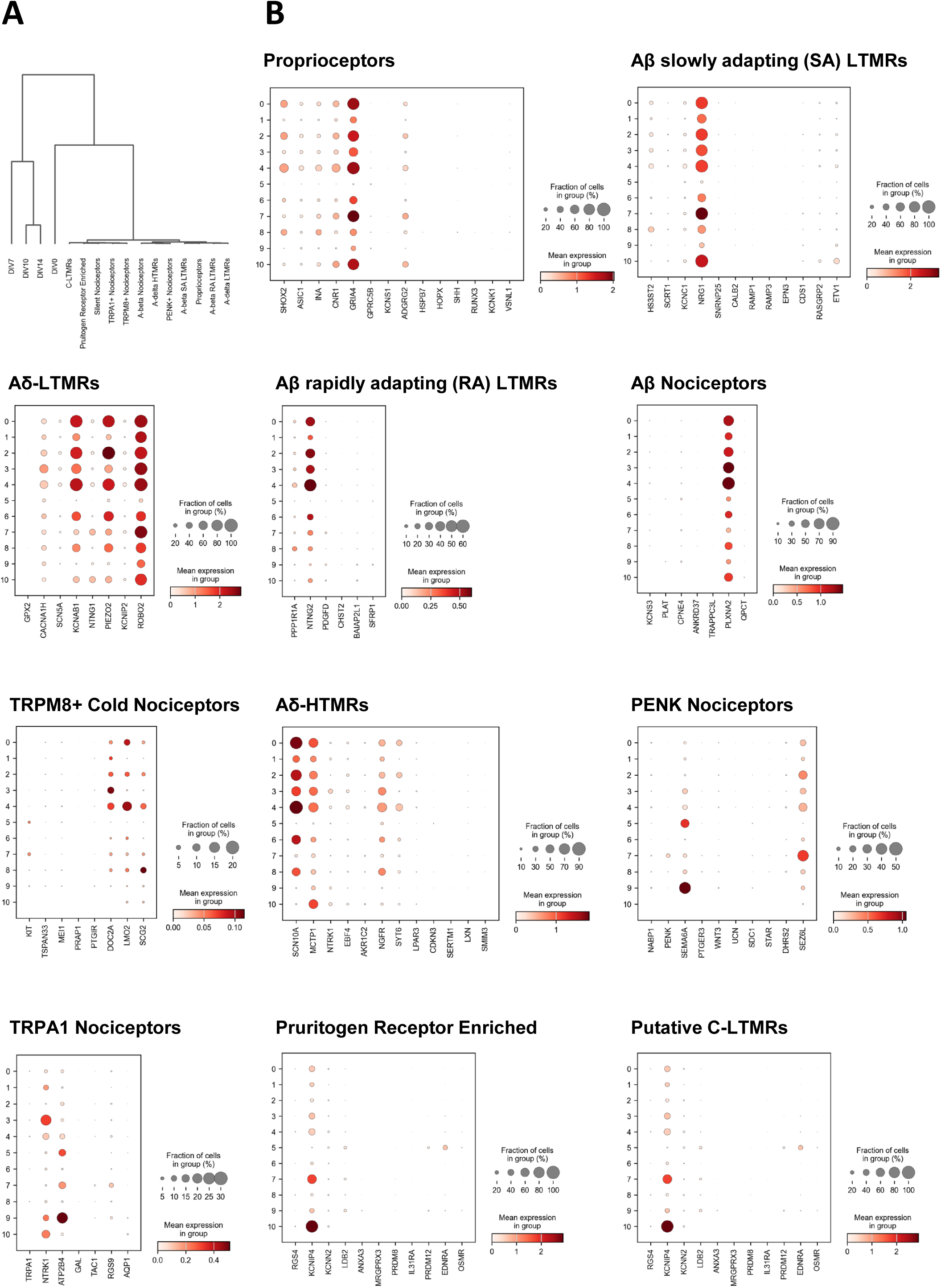
Single nucleus RNA sequencing of RealDRG reveal expression of a range of mechanoreceptor and nociceptor marker genes. **A)** Dendrogram clustering of RealDRG at DIV0, DIV7, DIV10, DIV14 with previously identified nociceptor subtypes. **B)** Dotplot expression of nociceptor subtype marker genes across Leiden clusters.

### 6. Functional characterization of RealDRG using automated patch clamp

Recent work has demonstrated the feasibility of applying automated high-throughput patch clamp techniques to study voltage-gated sodium (Na_v_) channels in native rodent DRG neurons [43] and RealDRG [16]. In order to unlock the full potential of this technique for translatable, high-throughput pain drug discovery, we looked to improve previously published success rates and expand the range of ion channel targets that can be functionally detected in RealDRG neurons over a course of maturation using three automated patch clamp systems: the lower-throughput, Port-a-Patch, the medium-throughput Patchliner, and the high-throughput SyncroPatch 384 devices. Using an optimized cell dissociation protocol, we were able to obtain success rates ranging between 35-60% for cells matured over 14-28 days in vitro (DIV) (**Figure 6A**; **Table 1**). Inward Na_V_ (including TTX-sensitive and TTX-resistant components), outward K_v_ currents and functional P2X, Piezo and GABA_A_ mediated-responses were seen in 80-100% of cells, while action potential firing increased from 40 to 80% of cells over 3 weeks (**Figure 6A**; **Table 1**). Seal quality was excellent (>300 MΩ, average) demonstrating high throughput sampling was feasible from over 100 cells per run, enabling comprehensive statistical analysis of these sensory neurons in less than an hour.

**Figure 6.**
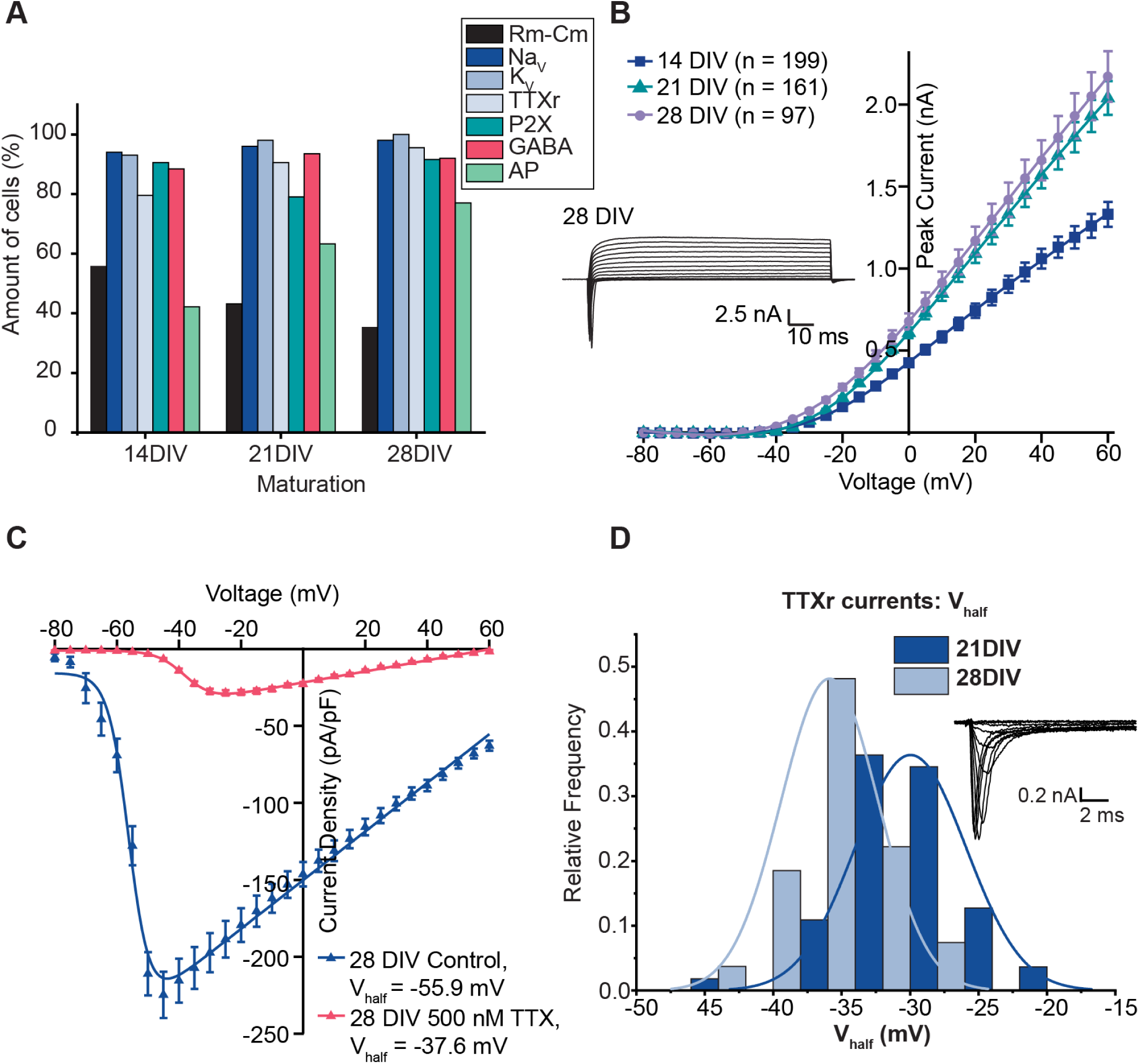
Voltage-gated currents recorded from RealDRG sensory neurons using APC. **A.** Summary of experimental success rates for cell capture, cells with K_V_, Na_V_ currents, P2X, GABA_A_ ligand-gated ionotropic receptor responses, and action potentials (AP). There appear to be some changes over time. **B.** K_V_ current-voltage plot at 14, 21 DIV and 28 DIV recorded on the SyncroPatch 384 with corresponding traces from an exemplar cell at 28 DIV. **C.** NaV current density at DIV 28 in the absence (dark blue) and presence (red) of TTX. The IV curve shifts to the right and the V_half_ more positive in the presence of TTX, consistent with Na_V_1.8. **D**. Histogram of V_half_ values calculated from IV recordings of TTXr Na_V_ channels at 21 DIV in dark blue and 28 DIV in light blue (recorded on the SyncroPatch 384). There is a shift to more negative potentials over time. Example traces recorded at 28 DIV are shown in the inset.

**Table 1:** Electrophysiological parameters for RealDRG at different maturation times (14, 21 or 28 DIV) recorded on the SyncroPatch 384. Current density for Na_V_ was recorded at 0 mV; TTXr was recorded at 10 mV; K_V_ was recorded at 40 mV; P2X-mediated currents were recorded at a holding potential of -80 mV and were responses to 20 µM for 1 s; and GABA-mediated currents were recorded at a holding potential of -80 mV and were responses to 100 µM GABA for 1 s. Number of cells for each parameter is given in parentheses.

| Parameter | 14 DIV | 21 DIV | 28 DIV |
| --- | --- | --- | --- |
| Seal Resistance ($\text{G}\Omega$) | $1.1 \pm 0.07$ (214) | $0.96 \pm 0.1$ (164) | $1.0 \pm 0.1$ (97) |
| Cell Capacitance (pF) | $8.0 \pm 0.5$ (220) | $12.0 \pm 0.6$ (168) | $16.3 \pm 2.9$ (103) |
| $\text{Na}_V$ (pA/pF) | $314 \pm 15$ (202) | $314 \pm 16$ (160) | $235 \pm 20$ (235) |
| $\text{TTX}_r$ (pA/pF) | $30.2 \pm 3.4$ (155) | $33.6 \pm 1.7$ (143) | $28.4 \pm 1.7$ (89) |
| $\text{K}_V$ | $144 \pm 4.8$ (202) | $159 \pm 5.6$ (160) | $151 \pm 6.9$ (97) |
| P2X | $83.8 \pm 5.4$ (94) | $49.2 \pm 2.9$ (97) | $47.4 \pm 3.4$ (65) |
| $\text{GABA}_A$ | $130 \pm 9$ (136) | $121 \pm 12$ (100) | $207 \pm 49$ (13) |

### 7. Voltage-gated ion channels recorded from RealDRG using automated patch clamp

DRG sensory neurons express a wide range of voltage-gated inward Na_V_ and Ca_V_, and outward K_V_ currents that control pain transmission, which can be accurately measured in voltage clamp mode using APC (**Figure 6B-C**). TTX-sensitive Na_V_1.7 (SCN9A), and TTX-resistant Na_V_1.8 (SCN10A) and Na_V_1.9 (SCN11A) channels are of particular interest for analgesia, as mutations in these targets are associated with various pain states in human patients [44]. RealDRG sensory neurons express SCN9A and SCN10A RNA and protein, with lower levels of SCN11A, and voltage clamp recordings on the SyncroPatch 384, Patchliner and Port-a-Patch show large TTX-sensitive currents activating at negative potentials, and a significant (∼10 - 20%) component of TTX-resistant inward current that activates at more positive potentials, characteristic of Na_V_1.8 channels and matching previously published results [26] (**Figure 6C-D**). The kinetics of TTX-resistant currents in RealDRG iPSC sensory neurons, and their relative contribution to macroscopic I_Na_, are similar to those recorded from adult human DRG [45] but differ from that in rat DRG neurons, illustrating another advantage of recording from native cells on APC to elucidate key species-specific differences in the responses. The ability to examine these neurons using current clamp to measure action potential firing while assessing changes in excitability complements the voltage clamp analysis of the distinct currents expressed.

### 8. Ligand-gated ion channels recorded from RealDRG

To further demonstrate translational utility of hiPSC-derived nociceptors, we investigated the pharmacological responses of various ligand gated ionotropic targets that were highly expressed in RealDRG neurons. Due to the ability to test thousands of activators or inhibitors with the robotic liquid handler, we decided to focus on two well-known targets, P2X and GABA_A_ channels. (**Figure 7**).

**Figure 7.**
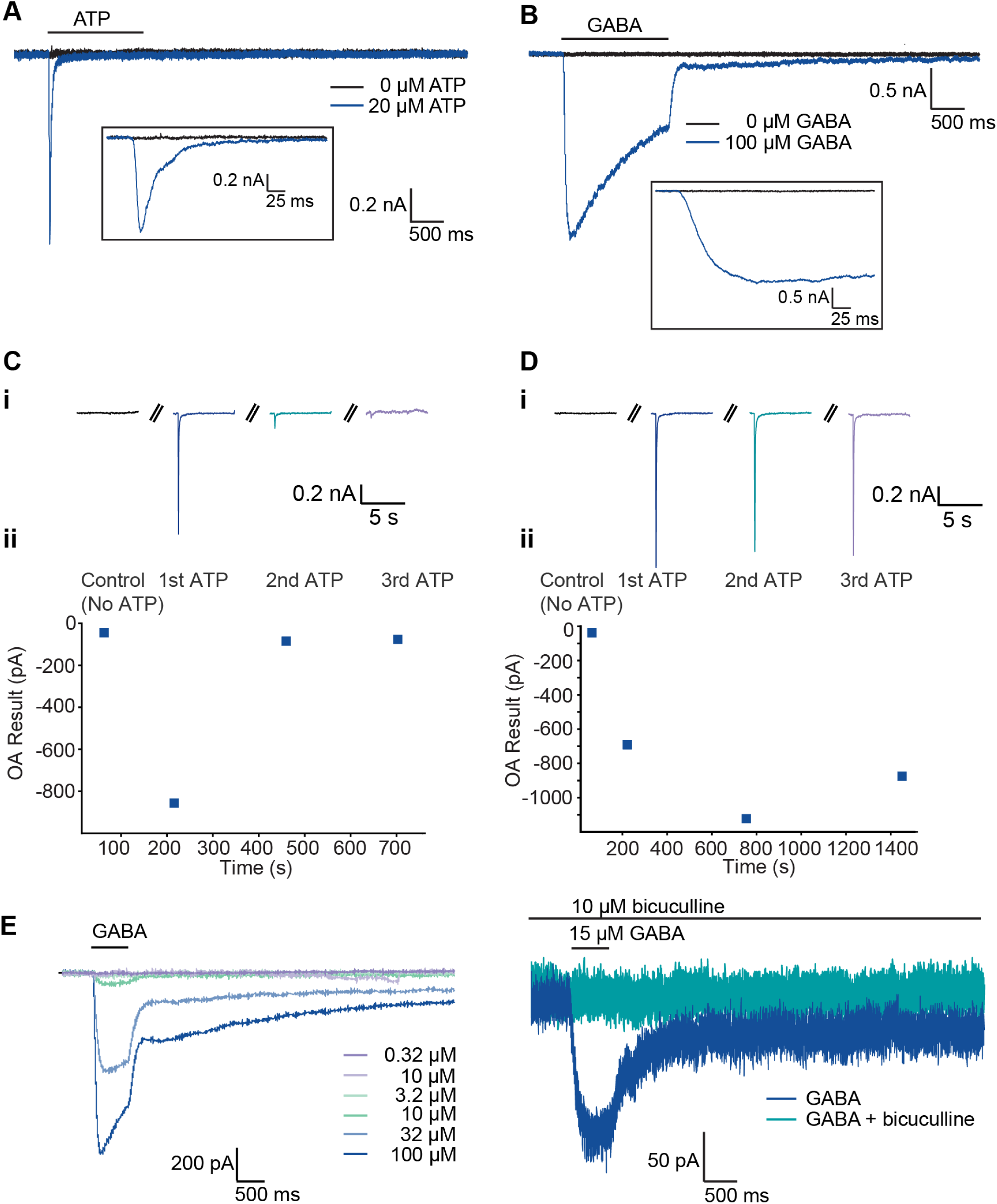
Ligand-gated ionotropic receptor currents. **A)** P2X receptor currents activated using 20 µM ATP on the SyncroPatch 384. Rapid activation and desensitization of ATP-evoked currents; inset at expanded time scale. **B)** GABA_A_ receptors activated by fast application of 100 µM GABA on the SyncroPatch 384. Inset shows time course of activation on an expanded time scale. **C**) Desensitization of P2X currents evoked every 4 min. This can be reduced to give consistent repeated responses in the same cell if ATP applications are made every 10 minutes, or (**D**) more rapidly in the presence of hexokinase to prevent ATP degradation. **E)** Currents activated by GABA were inhibited by the competitive GABA_A_ antagonist bicuculline (10 mM).

P2X_3_ receptor antagonists are promising drugs for the treatment of refractory chronic cough [46]. There are major differences in gene expression of P2X receptors between rodent and human DRG [47]. On the SyncroPatch 384 and Patchliner, we observed the expected rapid activation and inactivation of inward currents associated with P2X_3_-mediated currents (**Figure 7A**). In addition, we observed profound desensitization to repeated stimuli with ATP (**Figure 7C**). This effect could be effectively removed by extending the time between stimuli in the presence of hexokinase in the external solution, resulting in a stable response to repeated ligand application suitable for drug screening (**Figure 7D**).

GABA receptors are broadly expressed throughout the nervous system and have been associated with a wide variety of psychiatric and neurodevelopmental disorders including neuropathic pain [48]. Although DRG neurons lack intra-ganglionic synapses, GABA is a local neurotransmitter in this system. GABA_A_ Cl-channels are expressed in adult rat [49] and human DRG neurons [25], and their activation opens inward Cl-currents that hyperpolarize DRG neurons. GABA_A_ receptors can also be targeted by a wide variety of pharmacological compounds including benzodiazepines and general anesthetics. RealDRG express GABA related genes GABRA2, GABRA3, GABRB3, GABBR1, GABBR2 and we successfully recorded currents in response to application of 100 mM GABA that display the expected activation and inactivation kinetics (**Figure 7B**). As with the P2X recordings, GABA responses were detected in ∼90% of RealDRG neurons with current densities in the range of 125 – 200 pA/pF for GABA at 100 mM (**Figures 6A and 7B**). The response to GABA was concentration-dependent (**Figure 7E**) and blocked by bicuculline (**Figure 7F**), confirming that the responses are mediated by GABA_A_ receptors, although the subunit combination was not investigated.

### 9. Mechanoactivated currents in RealDRG are likely to be mediated by Piezo1 and Piezo2

Sensory neurons are highly sensitive to mechanical forces and are important for processing touch, proprioception and mechanical pain. Piezo ion channels are non-selective cation channels that transduce mechanical stimuli into electrical signals [50]. These channels open rapidly in response to membrane stretch and undergo rapid inactivation during continued application of the mechanical stimulus [51]. Piezo2 channels are thought to be the primary transducer of mechanical touch sensation in sensory neurons and expression of Piezo2 mRNA and functional channels have been demonstrated in RealDRG [17, 18]. While Piezo1 channels have been mostly implicated in mechano-sensation in non-neuronal cell types such as red blood cells and mRNA expression is lower in RealDRG neurons, there have been reports of Piezo1 expression and activity in DRG neurons [52, 53]). Recently a new technique, M-Stim, for high-throughput measurements of Piezo channel activity was developed in which fluid flow shear stress is applied acutely to cells under patch clamp and applied to Piezo1 channels in HEK cells [54]. We sought to extend the M-Stim technique to evaluate the presence of mechanically gated channels in RealDRG sensory neurons. Briefly, the automated liquid handling robot was programmed to deliver brief pulses of shear stress (20 µL at 100 µL/s) repeatedly. The standard protocol involves 3 rounds of stimulation with external buffer alone, a fourth application of buffer plus the Piezo1 selective potentiator, Yoda1, and a fifth application of buffer with GdCl_3_ as a control blocker of Piezo channels (**Figure 8 A,B**). Remarkably, 65% of all successfully patched cells survived all 5 rounds of M-Stim shear stress representing an overall success rate of 37%. Among these cells that survived the entire M-Stim testing paradigm, 31% had rapidly activating and inactivating inward currents of 50 pA or greater that are blocked by Gd^3+^. **Figure 8C** displays representative traces from 3 RealDRG neurons along with 3 Piezo1 expressing HEK cells highlighting the similarities in activation and inactivation kinetics. A larger proportion of cells (54%) responded to M-Stim in the presence of Yoda-1 which suggests the presence of functional Piezo1 channels in these cells. On average, the current amplitude in the presence of 10 mM Yoda1 was ∼80% larger than in the absence (**Figure 8Di**). In contrast, M-Stim measurements of Piezo1 channels in heterologous HEK expression system showed 400-800% larger currents in the presence of Yoda1 indicating that Piezo1 channels are responsible for only a fraction of the total current in RealDRG neurons and consistent with the possibility that Piezo2 channels are the major contributor to mechano-gated ionic currents in RealDRG neurons, which is supported by the high levels of Piezo2 expression. Lastly, perusal of individual cell responses to M-Stim revealed a variety of responses, shown in **Figure 8Dii-iv**. Responses ranged from stable with no modulation by Yoda1 (**Figure 8Dii**), potentiation over time with no apparent modulation by Yoda1 (**Figure 8Diii**), and cells which showed large potentiation by Yoda1 (**Figure 8Div**).

**Figure 8.**
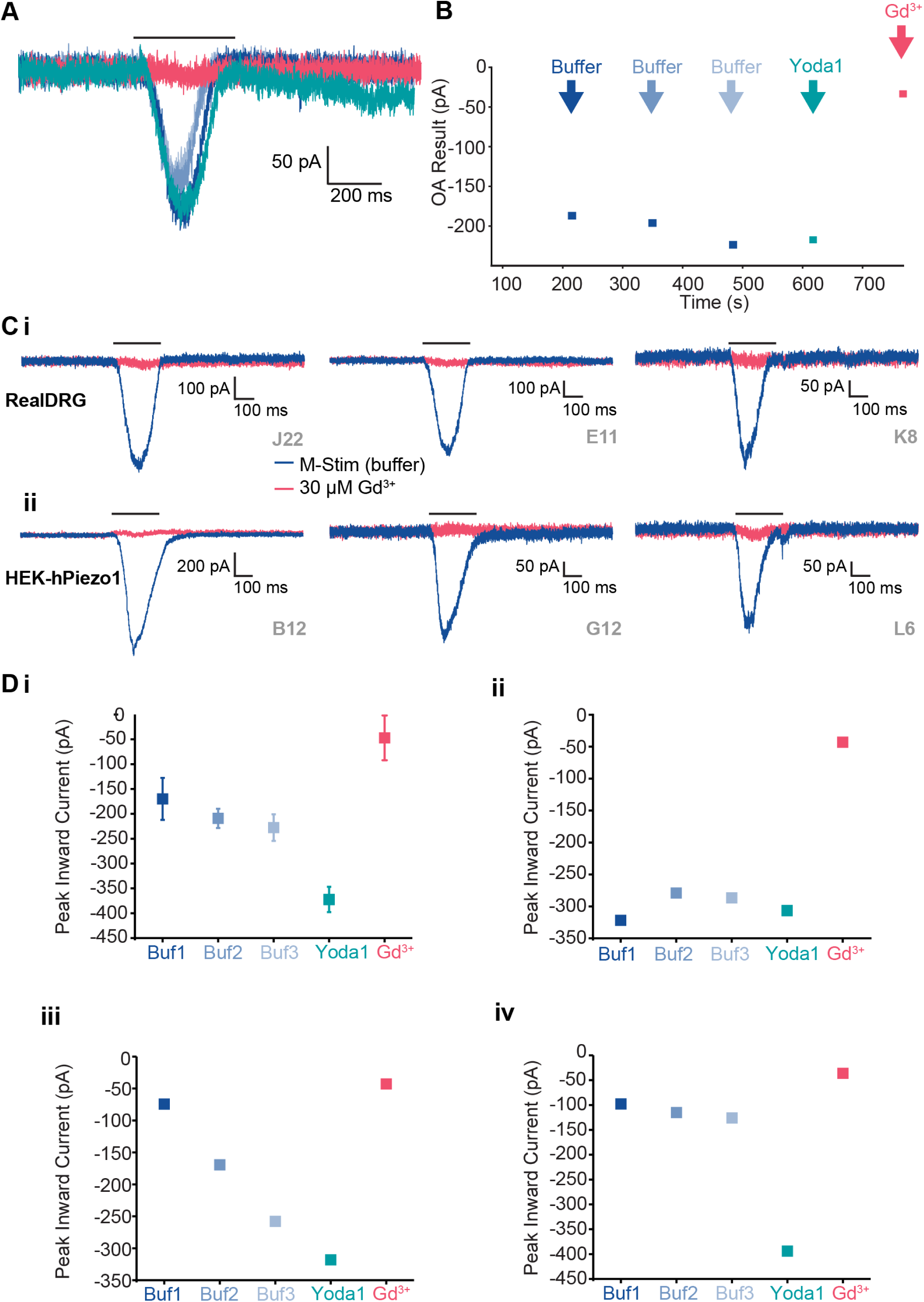
Mechano-gated ionic currents in RealDRG. **A)** Current traces recorded from an exemplar RealDRG on the SyncroPatch 384 shown overlaid where blue traces are buffer-only, green trace is with Yoda1 and red trace is with GdCl_3_. **B)** Timecourse of the experiment indicating activation using mechanical stimulation (M-Stim) where buffer solution was pipetting 110 mL/s at -80 mV. This was repeated 3 times followed by a 4^th^ application containing Yoda1 at 10 mM and a final application containing GdCl_3_ at 30 µM. **Ci)** Sample current traces from Real DRG neurons and comparison with **ii.** HEK cells overexpressing hPiezo1 in response to M-Stim protocol. Blue traces are currents in response to M-Stim with buffer only and red traces are currents in response to M-Stim in the presence of 30 mM GdCl_3_. **D)** Plot of peak inward current over time during M-Stim protocol. **Di** represents average ± SEM from all cells with current >50 pA in response to buffer alone. **ii – iv** are representative examples of individual neurons highlighting various response profiles: **ii** shows stable responses with no Yoda1 effect, **iii** shows current sensitization over time with no clear Yoda1 effect and **iv** shows stable responses to buffer followed by Yoda1 potentiation. All currents were blocked by GdCl_3_.

### 10. Action potential firing in RealDRG neurons

Following a detailed characterization of voltage-gated, ligand-gated and mechano-activated currents in voltage clamp mode, we investigated whether these cells could fire action potentials in current clamp mode using automated patch clamp. In a small number of RealDRG, spontaneous action potentials were recorded (**Figure 9A**), although these were only observed in 3.3% or 3.5% of cells at 21 and 28 DIV, respectively. In order to elicit action potentials, cells were held with holding current to maintain a resting membrane potential of -90 mV and then a staircase current protocol shown in **Figure 9B** was used to investigate the threshold at which action potentials were elicited in each cell. **Figure 9C** shows several examples of different firing profiles in RealDRG on the SyncroPatch 384, suggestive of possible functional heterogeneity within the population, potentially related to differences in cell identity or maturity. We observed similar action potential firing patterns regardless of APC platform used (**Figure 9, Supplementary Figure 3**). Complex ramp and step protocols can be utilized to assess sensitive changes in membrane excitability which are known to be involved in pathological or pain signaling. This enables proper categorization of single sensory neurons based on their action potential shape as well as their ability to respond to pharmacological agents. We used one NaV blocker, TTX, and could completely abolish action potential firing in RealDRG upon application of TTX, whilst AP firing in vehicle remained stable (**Figure 10**).

**Figure 9.**
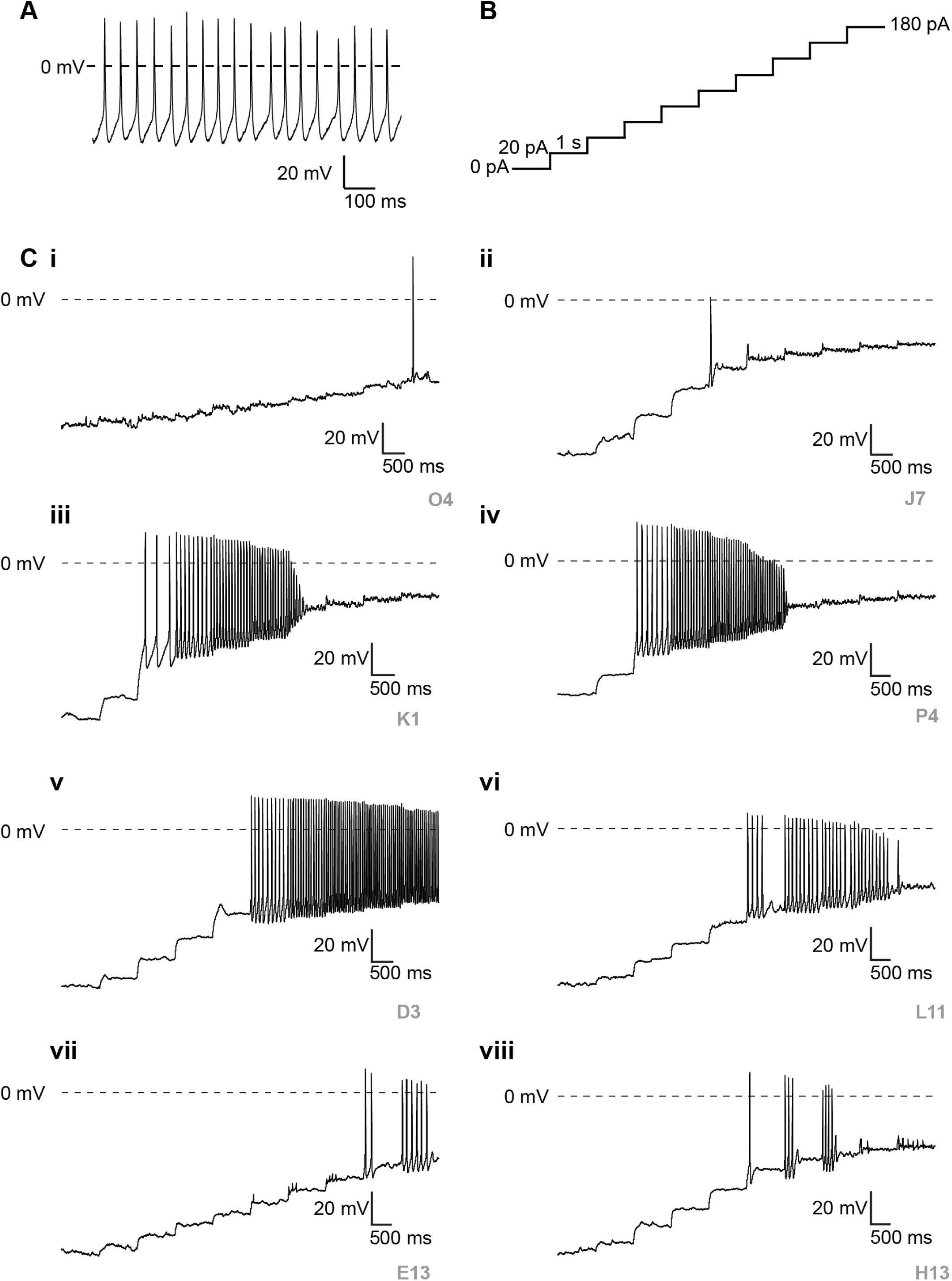
Spontaneous and elicited action potentials in RealDRG. **A.** Spontaneous action potentials were observed in a small subset of RealDRG neurons when no current was injected on both SyncroPatch 384 and Patchliner (example shown recorded on the Patchliner, RMP approx. -45 mV). **B**. Staircase protocol used to elicit action potentials on the SyncroPatch 384. Injected current was increased by 20 pA in 1 s steps to find the threshold for eliciting action potentials in each cell. **C.** Different action potential profiles for individual cells, ranging from a single action potential at a given current (**i** & **ii**), bursts of action potential firing (**iii** – **v**), and intermittent action potential bursts (**vi** – **viii**).

**Figure 10.**
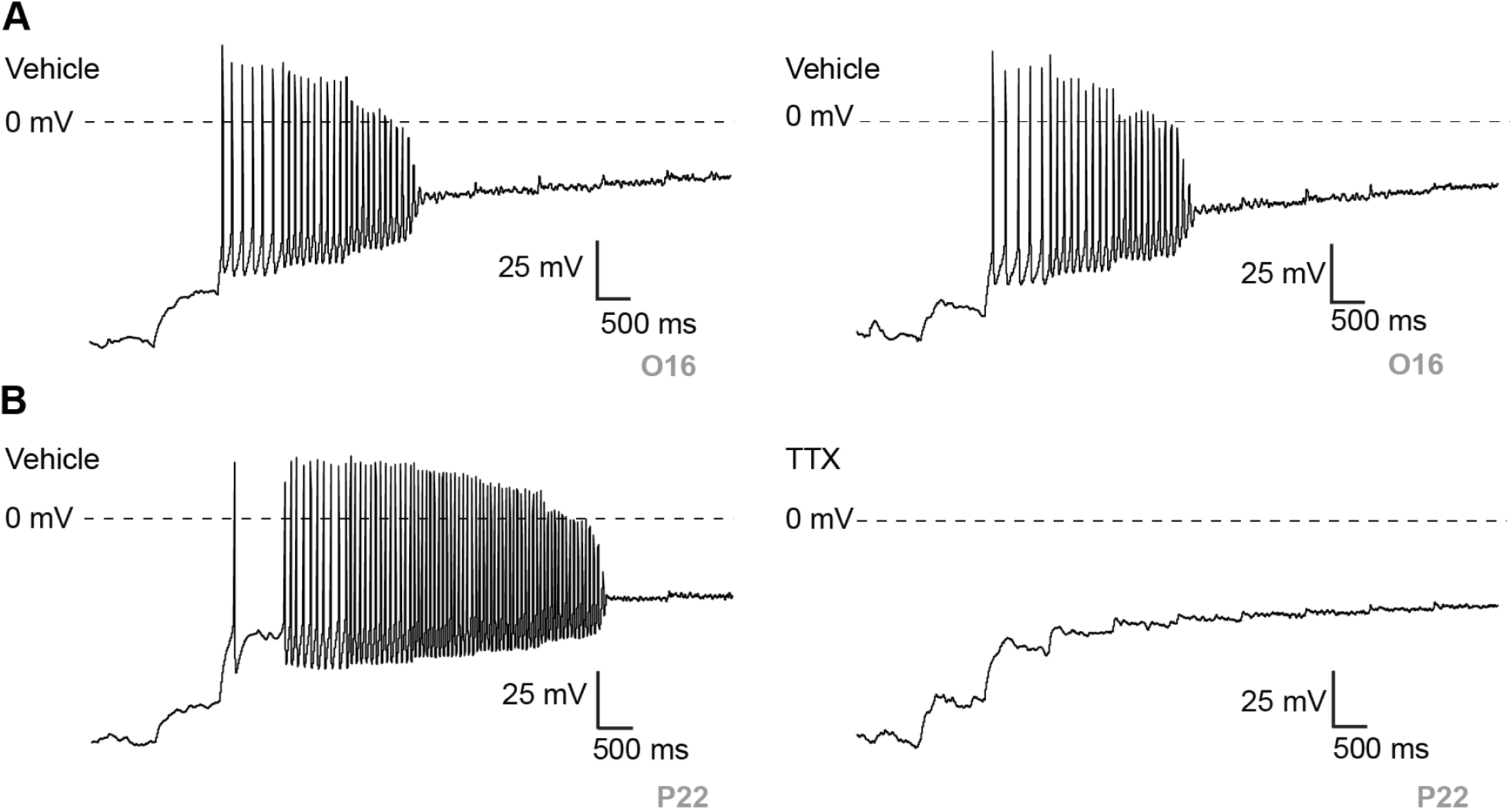
Elicited action potentials in RealDRG are abolished by TTX. **A.** Action potentials were elicited using the protocol shown in Figure 9B in vehicle (**A**) and in the presence of TTX (**B**) after incubation. TTX at completely abolished action potentials in RealDRG. Data recorded on the SyncroPatch 384.

## Discussion

This comparative study provides a number of new molecular insights into hiPSC-derived sensory neurons. No current hiPSC-derived sensory neuron completely recapitulates primary hDRG tissue on a bulk level, as hDRG tissues also contain a number of other cell types like satellite glia, Schwann cells, and immune cells. These additional non-sensory cells are expected to dilute the total amount of RNA contributed by sensory neurons, highlighting the extremely high levels of nociceptive transcripts present in native hDRG sensory neurons in comparison to any hiPSC-derived sensory neuron. To benchmark hiPSC-derived sensory neurons against native hDRG, therefore, it would be expected the expression level for any transcript found in a pure population of hiPSC-derived sensory neurons would likely need to exceed the expression found in hDRG for the two populations to approach parity. Alternatively, bulk-sequenced hiPSC-derived sensory neurons could be benchmarked against pseudo-bulked single-cell transcriptional analyses of native hDRG to understand relative expression levels, and therefore parity. Given these caveats, it appears the data produced by RealDRG sets a new standard in terms of transcriptomic equivalency between hiPSC-derived sensory neurons and native hDRG. Furthermore, numerous nociceptive transcripts found in RealDRG have been shown to be functional by high throughput APC, demonstrating an expression-function relationship. Given the scalability of hPSC-derived systems, RealDRG provide an essential resource for drug discovery for targets that have been validated, such as those that appear in this publication.

With many hiPSC-derived cell types, time in culture is often used as a solution toward embryological maturation. The present findings suggest this to be true assuming a directed differentiation methodology is used, such as in Clark, Deng, and Anatomic. However, maturation timelines can be accelerated when using directed differentiation depending on the environment to which the cells are exposed. The Anatomic protocol drastically accelerates embryological progression into an immature sensory neuron, which then can be accelerated toward maturation through time in comparison to the other directed differentiation protocols. Future refinements to maturation post-differentiation will need to be made to further increase nociceptive transcripts, with the idea that the transcription level needs to be increased to parity with hDRG to demonstrate functionality. Refinements could come in the form of additional soluble factors present in the media that more closely mimics the in vivo environment, or the addition of supportive cell types in co-culture found in the peripheral sensory nervous system, such as satellite glia and Schwann cells. It is possible the innervation targets of sensory neurons play a role in their maturation, so the inclusion of peripheral targets such as keratinocytes or central targets such as dorsal horn neurons will need to be studied. Finally, the unique anatomy of sensory neurons may need to be recapitulated, possibly with microfluidic chips, to further increase fidelity to their in vivo counterparts. These new lines of inquiry have yet to be explored, and provide the greatest potential to improve present culture systems. It is becoming clear, however, that strategies involving exogenously overexpressed identity transcription factors are unlikely to be the solution to improved culture performance, given that they currently pale in comparison to existing methods reliant on directed differentiation. The present work finds that these populations, even after advanced chronological aging, appear to be transcriptionally “frozen in embryological time” at the neural crest stage and never approaching equivalence to adult human DRG. Given transcription precedes translation and function, this poor transcriptional equivalence bodes poorly for their functional relevance and utility in pain drug discovery. To summarize the two methodologies used to differentiate hiPSCs as it relates to maturation— for sensory neurons at least— it appears the embryological “journey” encoded by morphogens is more important than the “destination” transcription factors that are overexpressed.

While modulating expression levels in a model system can be useful for predicting therapeutic efficacy in modalities such as antisense oligonucleotide therapy, pain electrophysiologists require a functional model that translates physiologically with dynamic conductance measurements. Utilizing the most high-throughput tool available for patch clamp, we demonstrate that a number of pain-related targets, such as voltage-gated sodium and potassium channels, ligand-gated GABA_A_, P2X receptors, and even mechanosensitive Piezo channels can be functionally characterized and assayed. Furthermore, this allows for unlimited pharmacological testing of the various ion channels expressed over time and may be a technique that can be optimized to interrogate other novel pain targets of interest, such as TRP channels or even intracellular Ryanodine receptors [55]. Recent automated organellar electrophysiological techniques may also be utilized to examine additional intracellular conductances in RealDRG neurons [56, 57]. Indeed, low throughput automated patch clamp devices have already been successfully used to record TPC and TRPML channels in lysosomes isolated from cell lines stably expressing TPCN2 [58, 59] or TRPML1 [60] or from podocytes [61] and scaling up the assay to higher throughput APC systems is ongoing [62]. Combining high throughput APC with organellar recordings from native cell types such as RealDRG may provide an important assay.

In conclusion, integrating transcriptomics with functional electrophysiology offers a robust approach to explore the molecular and cellular mechanisms underlying pain and to identify novel analgesic targets. Our study underscores the potential of RealDRG hiPSC-derived sensory neurons as both a scalable and relevant human model. This could significantly advance the development of more effective and specific pain therapeutics, ultimately improving the quality of life for patients suffering from chronic pain.

## Materials and Methods

### Differentiation of hiPSC Sensory Neurons

As previously described [16], the female hiPSC line (ANAT001) was cultured under feeder-free chemically defined conditions using TeSR-E8 medium and vitronectin as a coating substrate. Cells were single cell passaged onto a defined matrix and cultures were fed daily with a scaled-up version of Anatomic’s Senso-DM to produce immature sensory neurons by day 7 post-induction. Day 7 cultures were dissociated and cryopreserved. Lot-specific metrics were recorded including yield, cell number per vial, viability, post-thaw recovery, post-thaw viability, post-thaw morphology, purity, and sterility. Criteria used to determine lots passing quality control included neuronal purity > 95%, verified cell number per vial, post-thaw viability > 70%, and sterility. Additional detail related to differentiation, media compositions, materials used, and bioprocessing steps are proprietary information of Anatomic.

### Bulk RNAseq Analysis

Three lots of RealDRG manufactured from the same hiPSC line were thawed in Senso-MM (Anatomic Incorporated, #1030) at 100,000 neurons/cm^2^ in T25 format. Total RNA was collected at weekly timepoints for each line through four weeks. Pellets were formed by swirling cultures to release the cultures as a sheet before collection and centrifugation at 300g for 5 minutes. Cells were lysed and RNA extracted according to the RNA isolation kit instructions (Qiagen, #74104). After all samples had been collected, sequencing libraries were generated using Illumina TruSeq Stranded mRNA Library Prep Kit. Equimolar quantities from each library were sequenced on a NextSeq 550 High Output Kit v2.5 (75 cycles) at approximately 30 million single-end reads per sample. Publicly available datasets were retrieved from the SRA using the SRA-Toolkit (https://hpc.nih.gov/apps/sratoolkit.html). When available, paired end-reads were retrieved. For samples from Ray *et al.*, FASTQ files were uploaded to an S3 bucket and verified with md5sum hashes before processing. All samples were processed using the nf-core RNA-Seq pipeline (v 3.11.2) [63, 64]). Nextflow (v22.10.6) [65] using the ‘--docker’ profile to execute the workflow. Release 109 of the Human GRCh38 primary assembly was retrieved from ftp.ensembl.org. Release 43 of Gencode annotations for the Human primary assembly were retrieved from gencodegenes.org. Reads were aligned with STAR (v2.7.10a) and quantified using RSEM (v.1.3.1). In total, over 42 billion reads passed QC, or the equivalent of over 100 NextSeq lanes. For each sample, reads passing quality filters were used to generate transcript annotations using StringTie (v2.2.1). The Gencode annotation gtf was used to guide transcript assembly. Next, all novel transcript assemblies were merged and compared to the Gencode annotation using gffcompare (v0.12.6). The transcript map (.tmap) file was processed in R and class ‘s’ transcript annotations were discarded. RSEM quantifications of both gene and transcript level counts were loaded using the tximeta (v1.16.1) package with R 4.2.2. Transcripts with minimal expression (< 5 counts across all samples) were discarded. The function DESeqDataSetFromTximport was used to import quantifications and create a DESeq2 (v1.38.3) object for differential transcript and gene expression analysis.

### Single Cell RNAseq Analysis

Cells were cultured according to methods to DIV0, DIV7, DIV10, and DIV14. Adhered sensory neurons were washed from cell culture plates with PBS and centrifuged at 500g for 5 minutes at 4°C to form a pellet. Following pellet aggregation, cells were lyzed and nuclei were isolated according to manufacture protocol for the Nuclei Isolation Kit: Nuclei EZ Prep (Sigma-Aldrich). Nuclei were then passed through a 40 µm strainer to remove cellular debris and fixed according to manufacturer protocol within the Parse Biosciences fixation kit. Fixed nuclei from each collection point were stored at -80°C in a Mr. Frosty Freezing Container (Thermo Scientific). After all samples had been collected, sequencing libraries were generated using the Evercode Whole Transcriptome 100k v1 Kit (Parse Biosciences). Each sample used 12 of the available 48 wells of the library preparation kit, and individual nuclei were distributed evenly among wells. The libraries were sequenced on a NextSeq 550 High Output Kit v2.5 (300 Cycles) with a single 6 bp index. Prior to sequencing, a 5% PhiX spike-in was added according to manufacturer suggestion. Demultiplexing of sequencing libraries was performed using the Parse Biosciences Pipeline (v1.0.5p) to generate count matrices for individual samples. All downstream analysis including quality control, dimensional reduction, leiden clustering, and differential gene expression was performed within the Scanpy package (v1.9.2).

### Automated patch clamp recordings

Cells were thawed and plated into T75 flasks coated with iMatrix 511-SILK at a seeding density of 40,000 cells per cm2 and matured in Senso-MM. Cells were harvested using the Worthington Papain Kit (Worthington Biochemical Corp., Lakewood, NJ, USA) with slight modifications to achieve gentle dissociation of the neurons. Briefly, the neurons were incubated with 5 mL/T-75 flask of the papain/DNAase solution for 75 minutes at 37C followed by gentle trituration until free floating individual soma were observed. This suspension was carefully layered onto 6.5 mL of ovomucoid inhibitor and centrifuged for 3.75 minutes at ∼190Xg. The cell pellet was resuspended with standard physiological external solution at densities between 50,000 – 80,000 cells/mL and stored in the CellHotel on the Patchliner (Nanion Technologies GmbH, Munich, Germany) or SyncroPatch 384 (Nanion Technologies GmbH, Munich, Germay) APC platform, before being dispensed into each well of the NPC-16 or NPC-384 chip, respectively. The standard physiological external solution for all automated patch experiments consisted of (in mM), 140 NaCl 4 KCl, 2 CaCl_2_, 1 MgCl_2_, 10 HEPES, 5 glucose with pH adjusted to 7.4 using NaOH and osmolarity in the range of 295-300 mOsm. The internal solution for measurements of action potentials and voltage-gate potassium channels consisted of (in mM), 110 KF, 10 KCl, 10 NaCl, 1.5 MgCl_2_, 2 Na-ATP, 10 EGTA, 10 Hepes with pH adjusted to 7.2 using KOH and osmolarity in the range of 282-288 mOsm. The internal solution for measurement of TTX-resistant NaV currents and P2XR currents consisted of (in mM), 110 CsF, 10 CsCl, 10 NaCl, 10 EGTA, 10 Hepes with pH adjusted to 7.2 with CsOH and osmolarity in the range of 282-288 mOsm. The internal solution for measurement of GABA receptor currents consisted of (in mM), 70 CsF, 60 CsCl, 10 EGTA, 10 Hepes, 2 Na-ATP with pH adjusted to 7.2 with CsOH and osmolarity in the range of 282-288 mOsm. Standard patch clamp electrophysiology internal (KF-based) and external solutions were used to record inward Na_V_ and outward K_V_ currents and action potentials, and NaV currents were isolated by using a CsF-based internal solution. Standard voltage and current clamp protocols were used to set holding potential and measure resting membrane potential, and voltage-gated currents and action potentials were evoked with step voltage or current pulses. Data was acquired using PatchControl 384 (Nanion Technologies GmbH, Munich, Germany) and analyzed using DataControl 384 (Nanion Technologies GmbH, Munich, Germany) on the SyncroPatch 384. Data on the Patchliner and Port-a-Patch was acquired using PatchMaster (HEKA Elektronic, Stuttgart, Germany) and analyzed using IgorPro (Wavemetrics, OR, USA).

## Data availability

Sequence data is available in GEO, accession numbers GSE275412 (bulk) and GSE275413 (single cell). A searchable version of the bulk RNAseq data is also made available at https://anatomicincorporated.shinyapps.io/realdrgene/.

## Acknowledgements

We would like to acknowledge Sanster Biotechnologies and Benjamin Gansemer for their advice and expertise regarding the bulk RNAseq analysis and Marc Rogers at Albion Drug Discovery Services Ltd for their electrophysiology advice and review of this manuscript.

---

**Supplementary Figure 1.**
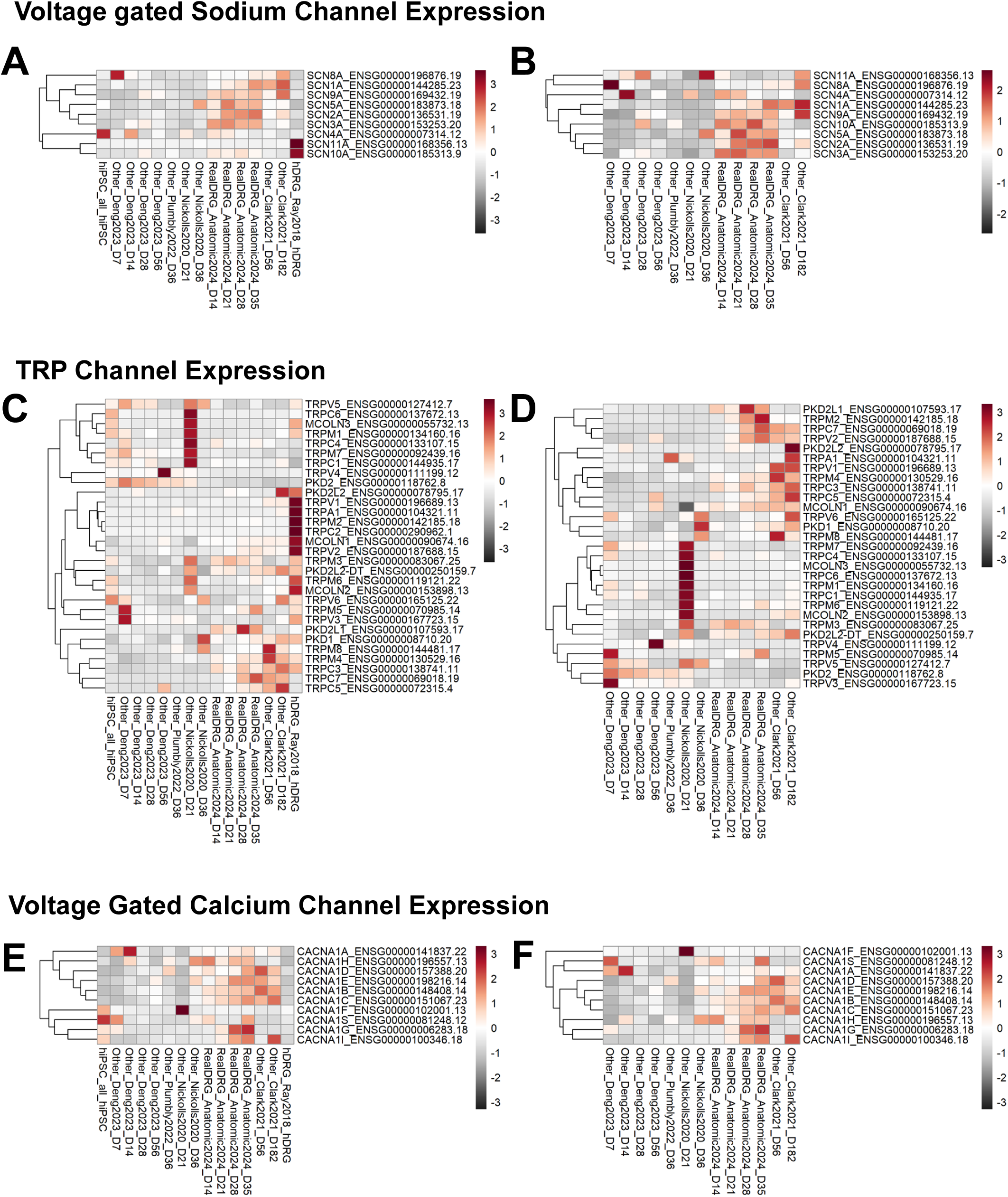
Voltage gated sodium channel expression across hiPSC derived sensory neuron populations **A)** with hDRG and **B)** without hDRG. TRP channel expression across hiPSC derived sensory neuron populations **C)** with hDRG and **D)** without hDRG. Voltage gated calcium channel expression across hiPSC derived sensory neuron populations **E)** with hDRG and **F)** without hDRG.

**Supplementary Figure 2.**
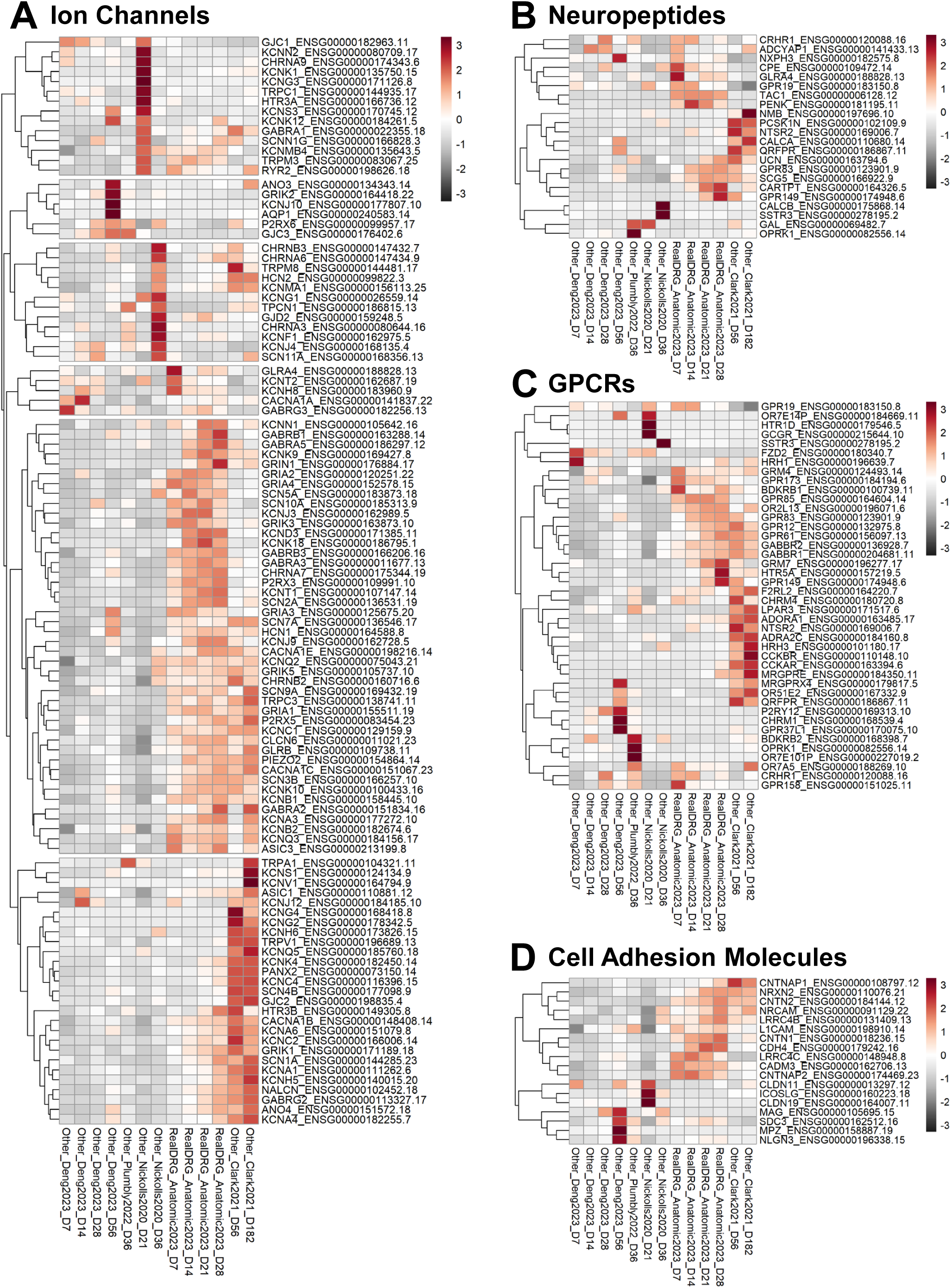
Heatmap analysis in only hiPSC-derived sensory neurons for specific **A)** Ion Channels **B)** Neuropeptides **C)** GPCRs and **D)** Cell adhesion molecules.

**Supplementary Figure 3.**
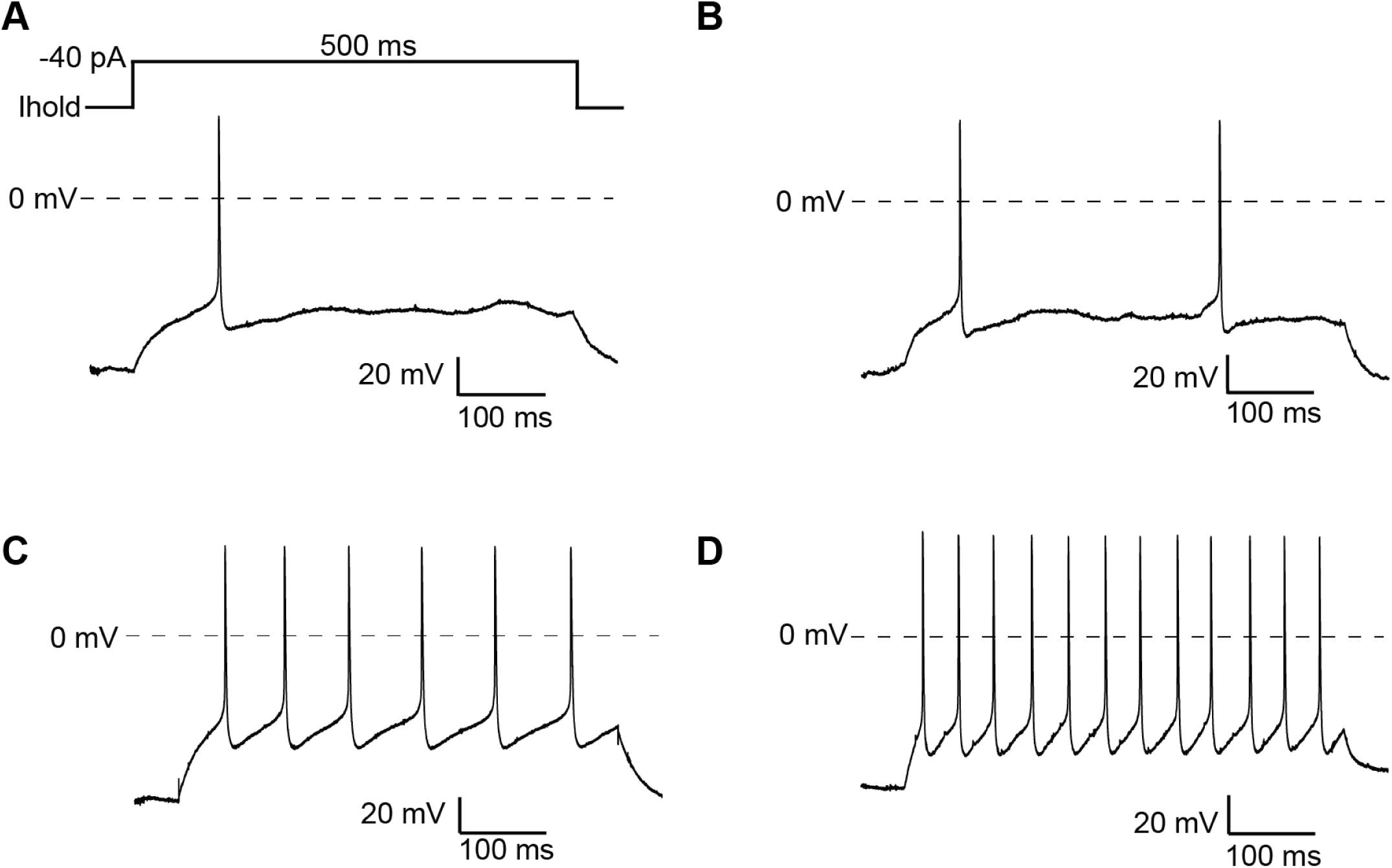
Elicited action potentials in RealDRG recorded on the Patchliner. Using the current protocol shown at the top of Panel A, action potentials were elicited in RealDRG on the Patchliner at 21 or 28 DIV. Similar to the data recorded on the SyncroPatch 384, different firing profiles were observed with a single or few action potentials elicited in some cells (**A, B**) or multiple action potentials elicited in others (**C, D**).

